# Genetic study links components of the autonomous nervous system to heart-rate profile during exercise

**DOI:** 10.1101/194167

**Authors:** Niek Verweij, Yordi van de Vegte, Pim van der Harst

## Abstract

Heart rate (HR) response to exercise, as defined by HR-increase upon exercise and HR-recovery after exercise, is an important predictor of mortality and believed to be modulated by the autonomic nervous system. However, the mechanistic basis underlying inter-individual differences remains to be elucidated. To investigate this, we performed a large-scale genome wide analysis of HR-increase and HR-recovery in 58,818 individuals. A total of 25 significant independent SNPs in 23 loci (P<8.3×10^−9^) were associated with HR-increase or HR-recovery, and 36 candidate causal genes were prioritized that were enriched for pathways related to neuron biology. There was no evidence of a causal relationship with mortality or cardiovascular diseases, however, a nominal association with parental lifespan was observed (5.5×10^−4^) that requires further study. In conclusion, our findings provide new biological and clinical insight into the mechanistic under-pinning of HR response to exercise, underscoring the role of the autonomous nervous system in HR-recovery.

**ABBREVIATIONS:**

BMI: Body mass index
ECG: Electrocardiography
HR: Heart rate
HRR: Heart rate recovery
GWAS: Genome-wide association study
LD: Linkage disequilibrium
MAF: Minor allele frequency
SE: Standard error
CI: Confidence interval

## INTRODUCTION

Physical activity places an increased demand on the cardiovascular capabilities of a person, it relies heavily on the regulation by the autonomic nervous system and cardiovascular health status^1^. Electrocardiograms (ECG) of exercise tests are used to determine cardiac fitness and function, and offers unique insights into cardiac physiology compared to rest ECGs. The first data linking electrocardiographic changes in response to exercise with mortality was presented in 1975, indicating that a low peak heart rate (HR) response during exercise was associated with an increased risk of cardiac death^2^. It is now well accepted that chronotropic incompetence confers a worse prognosis for cardiac mortality and events^3^. Increased HR during exercise, and HR-recovery after exercise is specifically associated with sudden cardiac death and all cause mortality, in healthy individuals^4–6^ and it is observed in coronary and heart failure patients regardless of β-blocker usage^7–9^. The majority of the studies were focused on HR-recovery at 60 seconds, which is heritable at around 60%^10^. The hypothesis linking HR-recovery to mortality arose from the work that associated components of the autonomic nervous system with sudden cardiac death^11^ and studies of decreased vagal activity^12,13^. McCrory et al.^14^ recently expanded on this topic by adding additional evidence, linking baroreceptor dysfunction with mortality. The study also showed HR-recovery in the first 10 seconds after an orthostatic challenge to be most predictive of mortality. The immediate response of the cardiovascular system to exercise is an increased HR that is attributable to a decrease in vagal tone, followed by an increase in sympathetic outflow and, to some extent circulating hormones^15^. The mechanism to reduce HR after exercise follows the inverse mechanism, a gradient of parasympathetic nervous system reactivation and sympathetic withdrawal^15^. The effect of this reactivation is believed to be strongest in the first 30 seconds after termination of exercise^16^. However, the exact molecular mechanisms underlying inter-individual differences in HR-response to exercise, as defined by HR-increase and HR-recovery, are unknown.

The UK Biobank includes a sub cohort of 96,567 participants that were invited for electrocardiographic exercise testing, which enabled for the first time in-depth genetic analyses of HR response to exercise. The aims of this study were (1.) to provide (shared) genetic heritability estimates among variables of the HR profile during exercise, (2.) to identify genetic determinants associated with HR-increase and HR-recovery at 10, 20, 30, 40 and 50 seconds and underlying candidate causal genes, and (3.) to obtain insights into pleiotropy and clinical consequences of HR-increase and HR-recovery.

## METHODS

### The UK Biobank cohort, ascertainment of the HR profile during exercise and quality control

UK Biobank is a cohort of individuals with an age range of 40-69 years that were registered with a general practitioner of the UK National Health Service. In total 503,325 individuals were included and provided informed consent between 2006 and 2010. The UK Biobank cohort was approved by the North West Multi-centre Research Ethics Committee. Detailed methods used by UK Biobank have been described elsewhere^17^.

In total 99,539 ECG-exercise records have been recorded in 96,567 participants that underwent a cardio assessment, 79,217 were recorded during the baseline visit (2006-2010), 20,322 at the second follow-up visit (2012-13). Participants were asked to sit on a stationary bike, start cycling after 15 seconds of rest and then perform 6minutes of physical activity, after which exercise was terminated and participants sat down for about 1 minute without cycling. The exercise protocol was adapted according to risk factors of the participant, details can be found elsewhere^18^. For this study, participants were only included if they were allowed to cycle at 50% or 30% of their maximum workload (no risk - minimum risk), as described further under “Statistical analyses (exclusions)”. The exercise was ended after reaching a pre-set maximum HR level of 75% of the age-predicted maximum HR. The Cardio assessment involved a 3 leads (lead I, II and III) ECG recording (AM-USB 6.5, Cardiosoft v6.51) at a frequency of 500 hertz. The ECG was recorded using 4 electrodes placed on the right and left antecubital fossa and wrist and stored in an xml-file of Cardiosoft.

Of all available ECG-records, 77,190 contained full disclosure data that could be used to detect R waves; others contained an error relating to the ECG device used (“*Error reading file C:\DOCUME∼1\UKBBUser\LOCALS∼1\Temp\ONL2F.tmp*”). R waves were detected with the gqrs algorithm^19^ and further processed using Construe^20^ (https://github.com/citiususc/construe) to detect individual Q-R-S waves. To obtain reliable RR intervals, following international recommendations ^21^, abnormal values (0,286 - 2 seconds) were removed. Additional outliers were removed by the tsclean function, part of R-package ‘forecast v7.3’ that incorporates the method described in Chen and Liu^22^ for automatic detection of outliers in time series. A total of 2,804 ECG's were excluded due to excess noise (identified by determining the standard deviation over a rolling standard deviation with a window length of 3 beats over RR intervals per ECG per phase and removing the 98^th^ percentile of this distribution). In total we inspected about 10 thousand RR interval profiles or ECGs to evaluate the RR-interval detection and quality control. For each ECG, we estimated the mean resting HR, the standard deviation of RR intervals (SDNN, log2 transformed) and the root mean square of successive differences between RR intervals (RMSSD, log2 transformed) from the RR intervals before exercise started. HR-increase was determined as the difference between peak HR during exercise and resting HR. HR recovery was defined as the difference between maximum HR during exercise and mean HR at 10±3, 20±3, 30±3, 40± and 50±3 seconds after exercise cessation (HRR10-HRR50). HR-recovery at exactly 1 minute was not available; only 9 participants recovered for a duration ≥60 seconds. Observations of the second follow-up visits were used when no baseline observation was available. Variables were inspected on normality and participants with extreme ECG exercise measurements (more than ±5 standard deviations from mean) were excluded on a per phenotype basis. By means of external validation, we estimated that resting-HR, SDNN and RMSSD were highly consistent with previous GWAS estimates^23,24^: β=1.085(se=0.029, P=3×10^−309^), β=1.145(se=0.051, P=1×10^−108^), and β= 1.0816(se=0.043, P=2×10^−139^), respectively as estimated by linear regression of the HR-phenotype on the polygenic score (described below). HR-phenotypes were rank-based inverse normal transformed to increase the power to detect low-frequency variants and to allow for comparisons of beta coefficients between traits. Source code, example data and further descriptions of the methods are available at https://github.com/niekverw/E-ECG.

Individual data on disease prevalence and incidence were derived from the Assessment Centre in-patient Health Episode Statistics (HES) and from self-reports during any of the visits through questionnaires and nurse-interviews as described previously^25^. Mothers, fathers and parental age of death was defined according to the study of Pilling et al.^26^, parental lifespan as a proxy for mortality was defined as the primary outcome variable.

### Genotyping and Imputation

Genotyping, quality control and imputation to 3 reference panels (HRC v1.1,1000 genome and UK10K) was performed by The Welcome Trust Centre for Human Genetics, as described in more detail elsewhere^27^. Sample outliers (based on heterozygosity or missingness) were excluded, as were 373 participants on the basis of gender mismatches. The analyses were restricted to HRC v1.1 SNPs with a MAF>1% and imputation quality score of >0.3.

### Statistical Analysis

#### Covariates

Regression analyses of resting-HR, SDNN and RMSSD were adjusted for gender, age, gender-age interaction, body mass index (BMI), BMI*BMI and the first 30 principal components and genotyping chip (Affymetrix UK Biobank Axiom or Affymetrix UK BiLEVE Axiom array). In addition, for HR increase and HR recovery the model also included exercise duration, exercise program (30 or 50% max load), maximum workload achieved and the interaction between the exercise program and maximum achieved workload to fully account for aerobic exercise capacity.

#### Exclusions

Participants were excluded if they stopped exercising earlier than planned, experienced chest-pain or other discomfort, were at medium to high cardiovascular risk^18^ at the time of the test, or terminated exercise due to unknown reasons. The population was stratified by participants that reported taking sotalol medication, beta-blockers, anti-depressants, atropine, glycosides or other anti-cholinergic drugs, or were diagnosed in history with a myocardial infarction, supraventricular tachycardia, bundle branch block, heart failure, cardiomyopathy or had a pacemaker or ICD implant in history. In a post-hoc sensitivity analysis the differences in beta estimates in these strata were assessed by a Chow test.

#### Analyses

In total 58,818 participants were included in the GWAS. GWAS and heritability analyses were performed using BOLT-LMM^28^ and BOLT-REML^29^ respectively, employing a conjugate gradient-based iterative framework for fast mixed-model computations that is able to accurately take into account population structure and relatedness, assuming additive effects. BOLT was modeled using 509,255 genotyped SNPs that were extracted from the final imputation set (to ensure a 100% call rate per SNP), and after pruning (*R*^2^ >0.5, using plink ‘--indep-pairwise 50 5 0.5); LD scores, also used by BOLT, were estimated from 2000 randomly selected UK Biobank participants (after sample exclusions based on relatedness, missingness and heterozygosity). To control for relatedness among participants in linear-logistic or cox- regression analyses, we used cluster-robust standard errors using genetic family ID as clusters. A family ID was given to individuals that were 3th-degree or closer based on the kinship matrix provide by UK Biobank (kinship coefficient >0.0442). All statistical analyses other than the genome-wide analysis were carried out using R v3.3.2 or STATA/SE release 13.

#### Statistical significance

Since this is by far the largest population based study of electrocardiographic exercise tests; independent cohorts that match this study in size and availability of variables (specific heart rate response variables and genetics) for replication are unavailable. Therefore, a conservative genome wide significant threshold of P<8.3×10^−9^ was adopted accounting for 6 independent traits, in accordance with similar multi-phenotype studies of this scale^30–34^.

#### Locus determination

Variants were considered to be independent if the pair-wise LD (r^2^) was less than 0.01. A locus was defined as a 1MB region surrounding the highest associated independent SNP. The strongest associated variant within a locus was assigned the ‘sentinel SNP’, however multiple ‘independent SNPs’ can be in one locus, which was confirmed by adjusting for the sentinel-SNP in the locus using linear regression.

### Pleiotropy analyses

The GWAS catalog database (https://www.ebi.ac.uk/gwas/) was queried by searching for SNPs in a 1MB distance of the SNPs found in this study. LD was determined by calculating the r^2^ and D’ in UK Biobank between the GWAS catalog SNPs and the SNPs found in this study.

To gain insights into pleiotropy among HR-variables, we performed linear regression analyses for all significantly associated SNPs with resting-HR, HR-variability (SDNN and RMSSD), HR-increase and HR-recovery and visualized the z-scores, aligned to the HR-recovery increasing allele.

### Polygenic score

Polygenic scores of HR-increase and HR-recovery were constructed by summing the number of alleles that increases HR-increase or HR-recovery, weighted (multiplied) for the corresponding beta-coefficients. The relationship between the polygenic score and clinical phenotypes were tested in 422,947 individuals that were not part of the discovery GWAS to avoid any potential bias or reverse confounding, using linear-, logistic- and cox -regression analyses. The statistical power for a case/control Mendelian randomization in UK Biobank (N=422,334) was calculated at alpha=0.05, as described previously^35^.

### Functional variants and candidate genes

To search for evidence of functional effects of SNPs at the HR-loci we used multiple QTL databases: STARNET (Stockholm-Tartu Atherosclerosis Reverse Network Engineering Task)^36^, GTEX version 6^37^, cis-eQTL datasets of Blood^38–40^ and cis-meQTLs^41^. Only eQTLS/meQTLs that achieved P<1x10^−6^ and were in LD (r^2^>0.8) with the queried GWAS SNP were considered significant. Multiple eQTLs were observed at the same SNPs in different tissues/studies, providing extra evidence for being a true eQTL.

Long range chromatin interaction in the 1 MB region at any of the sentinel SNPs were examined and visualized using HUGin^42^, only genes that showed a bonferonni significant association that showed a clear pattern of interaction between the sentinel SNP and the promoter region were prioritized.

Candidate genes were prioritized: a) by proximity - the nearest gene or any gene within 10kb, b) by protein-coding gene variants in LD (r^2^>0.8) with the sentinel SNP, c) by eQTLs (described above) and d) by long range chromatin interactions (described above).

## RESULTS

Participants of UK Biobank exercised for approximately 350 (±44.9) seconds, the mean duration of the recovery phase was 52.6 (±1.7) seconds; overall characteristics are presented in **Supplementary Table 1**. All HR phenotypes were normally distributed prior to rank-based inverse normal transformation.

To gain insights into the correlation among phenotypes of the HR profile during exercise, we first performed heritability analyses and genetic correlations across 9 HR phenotypes: the increase in HR from the resting level to the peak exercise level (HR increase); the decrease in rate from the peak exercise level to the level 10, 20, 30, 40 and 50 seconds after termination of exercise (HRR10-HRR50). Resting-HR and HR-variability as defined by SDNN an RMSSD were included for comparisons. The highest heritability estimates were observed for HR-recovery and HR-increase (h_2gSNP_=0.22). HR variability was much less heritable, h_2gSNP_ =0.12 and 0.14 for SDNN and RMSSD, based on SNP-heritability estimates by BOLT-REML (Fig. 1). All of the HR-variables were highly correlated with each other (Fig. 1), though HR-recovery and HR-increase were more correlated with each other (r=0.6 to 0.9) than with HR-variability (r=0.42 to 0.6) or resting-HR (r=-0.18 to -0.37). The genotypic correlations where slightly higher compared to the phenotypic correlations. All of the heritability and correlation estimates were highly significant (P<1×10^−8^).

**Figure 1.**
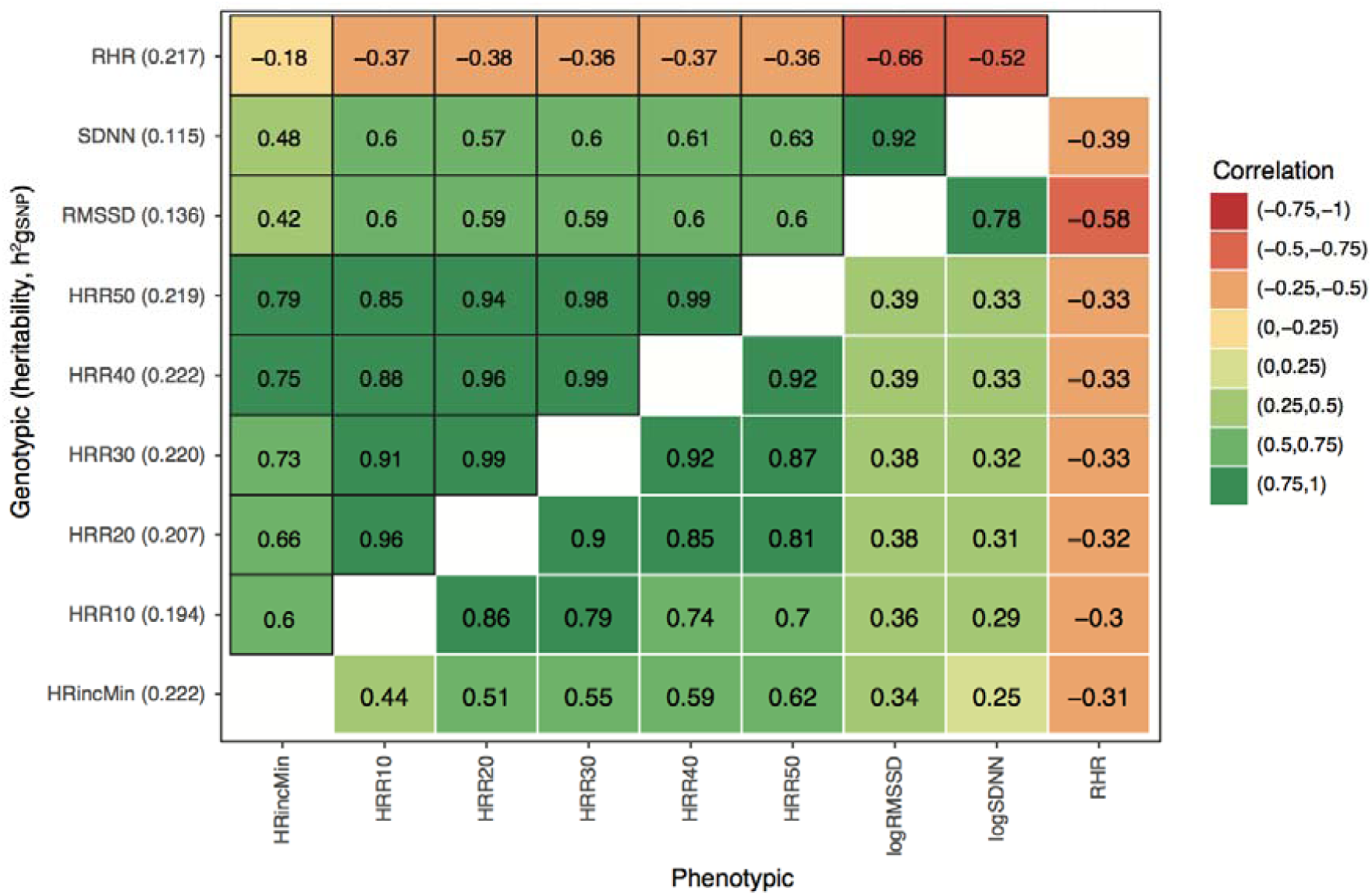
Genetic correlations (shared heritability), are shown above the diagonal, phenotypically observed correlations are below the diagonal. Heritability estimates of each trait are between brackets at the y-axis. All of the estimates shown here were highly significant (P<10^−8^). Correlations are based on the residual variance after adjustments for age, sex and BMI, exercise specific variables and genetic specific variables (only for the genetic correlations).

Genome wide association analyses were conducted of HR-increase and HRR10-HRR50. Twenty-three genomic loci defined by 1 MB at either side of the highest associated SNP were genome wide significant (Table 1, Fig. 2 **and Supplementary Table 2**), 2 additional independent signals in 2 loci were confirmed by conditional analyses (**Supplementary Table 3**). Rs6488162 in *SYT10* was the most significant genetic variant for all phenotypes (P=3.1×10^−30^ for HR-increase, to P=5.3×10^−66^ for HRR10). Results of the sensitivity analyses are shown in **Supplementary Table 4**, indicating that the SNP associations are not biased by participants receiving medication or having diagnoses of heart disease in history. **Supplementary Fig. 1** shows the regional association plots of each locus. To facilitate future studies we make available the methods and complete summary statistics of all genetic variants and traits at www.cardiomics.net and at open access repositories for GWAS summary statistics.

**Table 1.**
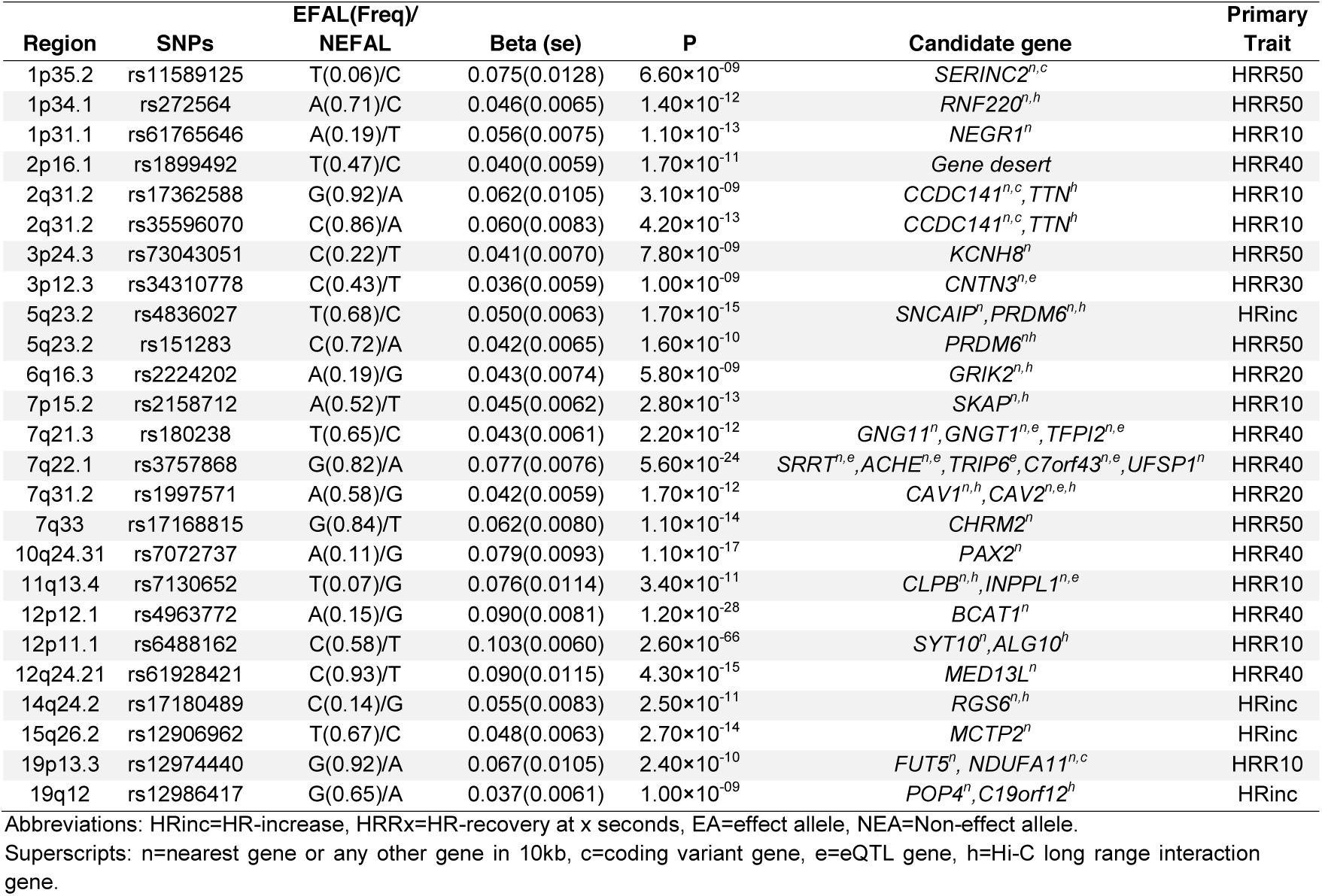
The GWAS identified 25 genome wide significant SNPs in 23 loci for HR-increase or HR-recovery. More detailed information can be found in **Supplementary Table 2** and **3**.

| Region | SNPs | EFAL(Freq)/<br>NEFAL | Beta (se) | P | Candidate gene | Primary<br>Trait |
| --- | --- | --- | --- | --- | --- | --- |
| 1p35.2 | rs11589125 | T(0.06)/C | 0.075(0.0128) | 6.60×10<sup>-09</sup> | *SERINC2<sup>n,c</sup>* | HRR50 |
| 1p34.1 | rs272564 | A(0.71)/C | 0.046(0.0065) | 1.40×10<sup>-12</sup> | *RNF220<sup>n,h</sup>* | HRR50 |
| 1p31.1 | rs61765646 | A(0.19)/T | 0.056(0.0075) | 1.10×10<sup>-13</sup> | *NEGR1<sup>n</sup>* | HRR10 |
| 2p16.1 | rs1899492 | T(0.47)/C | 0.040(0.0059) | 1.70×10<sup>-11</sup> | *Gene desert* | HRR40 |
| 2q31.2 | rs17362588 | G(0.92)/A | 0.062(0.0105) | 3.10×10<sup>-09</sup> | *CCDC141<sup>n,c</sup>, TTN<sup>h</sup>* | HRR10 |
| 2q31.2 | rs35596070 | C(0.86)/A | 0.060(0.0083) | 4.20×10<sup>-13</sup> | *CCDC141<sup>n,c</sup>, TTN<sup>h</sup>* | HRR10 |
| 3p24.3 | rs73043051 | C(0.22)/T | 0.041(0.0070) | 7.80×10<sup>-09</sup> | *KCNH8<sup>n</sup>* | HRR50 |
| 3p12.3 | rs34310778 | C(0.43)/T | 0.036(0.0059) | 1.00×10<sup>-09</sup> | *CNTN3<sup>n,e</sup>* | HRR30 |
| 5q23.2 | rs4836027 | T(0.68)/C | 0.050(0.0063) | 1.70×10<sup>-15</sup> | *SNCAIP<sup>n</sup>, PRDM6<sup>n,h</sup>* | HRinc |
| 5q23.2 | rs151283 | C(0.72)/A | 0.042(0.0065) | 1.60×10<sup>-10</sup> | *PRDM6<sup>nh</sup>* | HRR50 |
| 6q16.3 | rs2224202 | A(0.19)/G | 0.043(0.0074) | 5.80×10<sup>-09</sup> | *GRIK2<sup>n,h</sup>* | HRR20 |
| 7p15.2 | rs2158712 | A(0.52)/T | 0.045(0.0062) | 2.80×10<sup>-13</sup> | *SKAP<sup>n,h</sup>* | HRR10 |
| 7q21.3 | rs180238 | T(0.65)/C | 0.043(0.0061) | 2.20×10<sup>-12</sup> | *GNG11<sup>n</sup>, GNGT1<sup>n,e</sup>, TFPI2<sup>n,e</sup>* | HRR40 |
| 7q22.1 | rs3757868 | G(0.82)/A | 0.077(0.0076) | 5.60×10<sup>-24</sup> | *SRRT<sup>n,e</sup>, ACHE<sup>n,e</sup>, TRIP6<sup>e</sup>, C7orf43<sup>n,e</sup>, UFSP1<sup>n</sup>* | HRR40 |
| 7q31.2 | rs1997571 | A(0.58)/G | 0.042(0.0059) | 1.70×10<sup>-12</sup> | *CAV1<sup>n,h</sup>, CAV2<sup>n,e,h</sup>* | HRR20 |
| 7q33 | rs17168815 | G(0.84)/T | 0.062(0.0080) | 1.10×10<sup>-14</sup> | *CHRM2<sup>n</sup>* | HRR50 |
| 10q24.31 | rs7072737 | A(0.11)/G | 0.079(0.0093) | 1.10×10<sup>-17</sup> | *PAX2<sup>n</sup>* | HRR40 |
| 11q13.4 | rs7130652 | T(0.07)/G | 0.076(0.0114) | 3.40×10<sup>-11</sup> | *CLPB<sup>n,h</sup>, INPPL1<sup>n,e</sup>* | HRR10 |
| 12p12.1 | rs4963772 | A(0.15)/G | 0.090(0.0081) | 1.20×10<sup>-28</sup> | *BCAT1<sup>n</sup>* | HRR40 |
| 12p11.1 | rs6488162 | C(0.58)/T | 0.103(0.0060) | 2.60×10<sup>-66</sup> | *SYT10<sup>n</sup>, ALG10<sup>h</sup>* | HRR10 |
| 12q24.21 | rs61928421 | C(0.93)/T | 0.090(0.0115) | 4.30×10<sup>-15</sup> | *MED13L<sup>n</sup>* | HRR40 |
| 14q24.2 | rs17180489 | C(0.14)/G | 0.055(0.0083) | 2.50×10<sup>-11</sup> | *RGS6<sup>n,h</sup>* | HRinc |
| 15q26.2 | rs12906962 | T(0.67)/C | 0.048(0.0063) | 2.70×10<sup>-14</sup> | *MCTP2<sup>n</sup>* | HRinc |
| 19p13.3 | rs12974440 | G(0.92)/A | 0.067(0.0105) | 2.40×10<sup>-10</sup> | *FUT5<sup>n</sup>, NDUFA11<sup>n,c</sup>* | HRR10 |
| 19q12 | rs12986417 | G(0.65)/A | 0.037(0.0061) | 1.00×10<sup>-09</sup> | *POP4<sup>n</sup>, C19orf12<sup>h</sup>* | HRinc |

Abbreviations: HRinc=HR-increase, HRRx=HR-recovery at x seconds, EA=effect allele, NEA=Non-effect allele.
Superscripts: n=nearest gene or any other gene in 10kb, c=coding variant gene, e=eQTL gene, h=Hi-C long range interaction gene.

**Figure 2.**
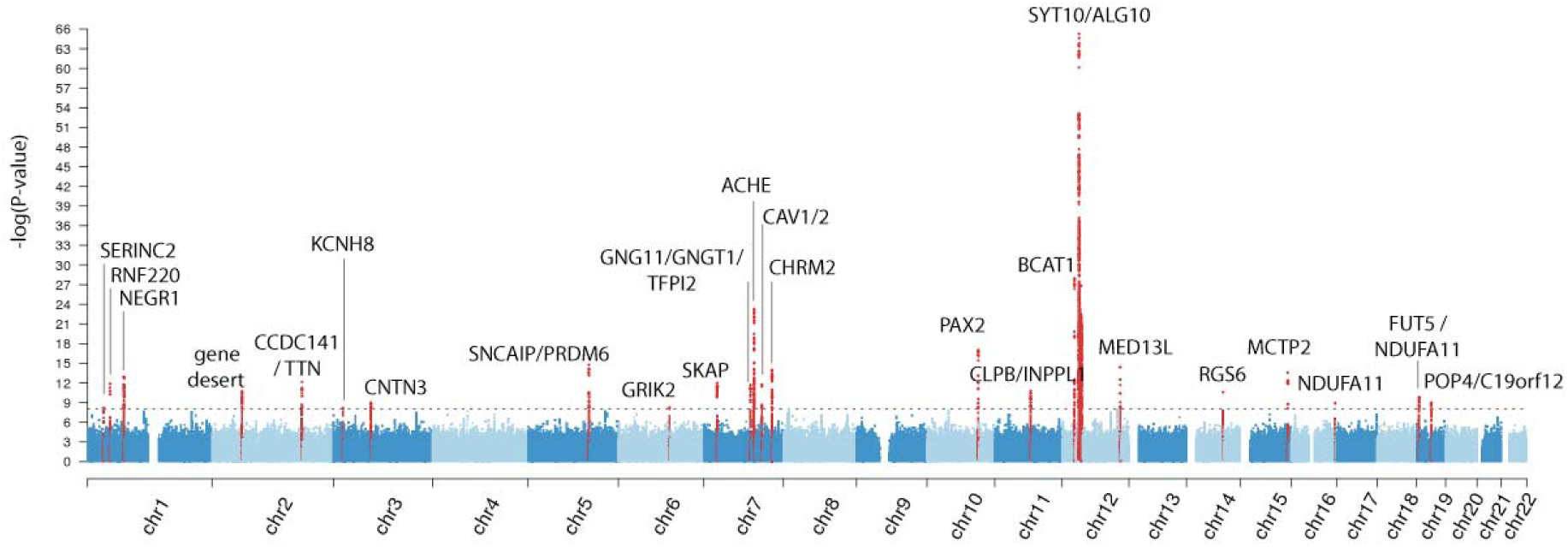
Manhattan plot of the GWAS of HR-increase and recovery, the smallest P-values per SNP across all of the 6 studied traits are shown, as depicted on the y axis, the x axis shows their chromosomal (chr) positions. Red dots represent genome wide significant loci (P<8.3×10^−9^).

### Insights into biology

A total of 36 candidate causal genes at the 23 loci were identified, 27 genes were prioritized by being near the sentinel SNP or within 10 kilobases, 3 genes were prioritized by coding variants in LD of r^2^>0.8 with a sentinel SNP (**Supplementary Table 5**), 10 genes were prioritized by eQTL analyses (**Supplementary Table 6**) and 11 genes were prioritized by long-range interaction analyses in Hi-C data (**Supplementary Table 7** and **Supplementary Fig. 3**). Multiple lines of evidence could prioritize a gene (shown in Table 1), which may be helpful for prioritizing the most likely candidate genes and mechanisms at each locus.

Pathway analyses were attempted with ‘DEPICT’^43^, a tool that prioritizes genes, pathways and tissues by using the genomic region surrounding SNPs as input, however there were no significant pathways or tissues after correcting for multiple testing. In turn, GeneNetwork (http://129.125.135.180:8080/GeneNetwork/pathway.html)^44^ was used, which employs the same underlying co-expression dataset (based on GEO data) but allows only the 36 candidate genes as input. The candidate genes were enriched for terms related to neurons and axons (‘axon guidance’, ‘neuron recognition’, ‘peripheral nervous system neuron development’, ‘synapse’), gap-junctions (‘adherens junction organization’ and ‘gap junction’), but also included 'catecholamine transport' and ‘decreased dopamine level’, among others (**Supplementary Table 8**). Nerve tissue was also highly enriched compared to other tissues in a separate analysis based on the GTEx dataset (P<0.01, **Supplementary Figure 4.**).

### Insights into pleiotropy and clinical relevancy

To increase our understanding of potentially mediating mechanisms at the genetic variant level we searched the literature and the GWAS-catalog for previously reported variants. Of the 23 loci, eleven are in high LD (r^2^>0.6) with previously identified SNPs for resting heart rate^24,45^ or heart rate variability^23^ (**Supplementary Table 9**). A wider search in the GWAS-catalog revealed that SNPs in high LD (r^2^>0.6) with rs61765646 (*NEGR1*) were reported for the association with obesity; rs17362588 (*CCDC141/TTN,* but not the independent SNP rs35596070) and rs12906962 (*MCTP2*) with diastolic blood pressure, rs7072737 (*PAX2*) with systolic blood pressure; rs4963772 (*BCAT1*) with PR-interval and rs1997571 (*CAV1*) with atrial fibrillation and PR interval (**Supplementary Table 10**). The majority, fifteen of 23 loci, has not been previously identified in any genome wide associated study.

Because a large proportion of the loci were already reported for their association with other HR-phenotypes, we examined SNP association with the different HR traits in the current study, in order to entangle the effects and identify SNPs that are primarily driven by HR-increase and HR-recovery. Linear regression analyses were performed across all associated SNPs and traits, and adjusted the associations for (1.) resting-HR, (2.) resting-HR and HR-variability (3.) resting-HR, HR-variability and HR-increase (**Supplementary Table 11** and visualized in Fig. 3). Fig. 3 illustrates rs17362588 (*TTN*/*CCDC141*) to be primarily associated with resting-HR and highlights the following loci for HR-variability: rs17180489 (*RGS6*), rs12974440 (*NDUFA11*) and to a lesser degree rs180238 (*GNG11, GNGT1, TFPI2*) as the associations with HR-recovery and HR-increase were diminished significantly upon extra adjustments of SDNN and RMSSD. The analyses also indicated that rs272564 (*RNF220*), rs4836027 (*SNCAIP/PRDM6*), rs4963772 (*BCAT1*), rs12906962 (*MCTP2*) and rs12986417 (POP4) were primarily associated with HR-increase following extra adjustments of HR-increase. In total sixteen SNPs remained independently associated with HR-recovery, including the most significant locus *SYT10*.

**Figure 3.**
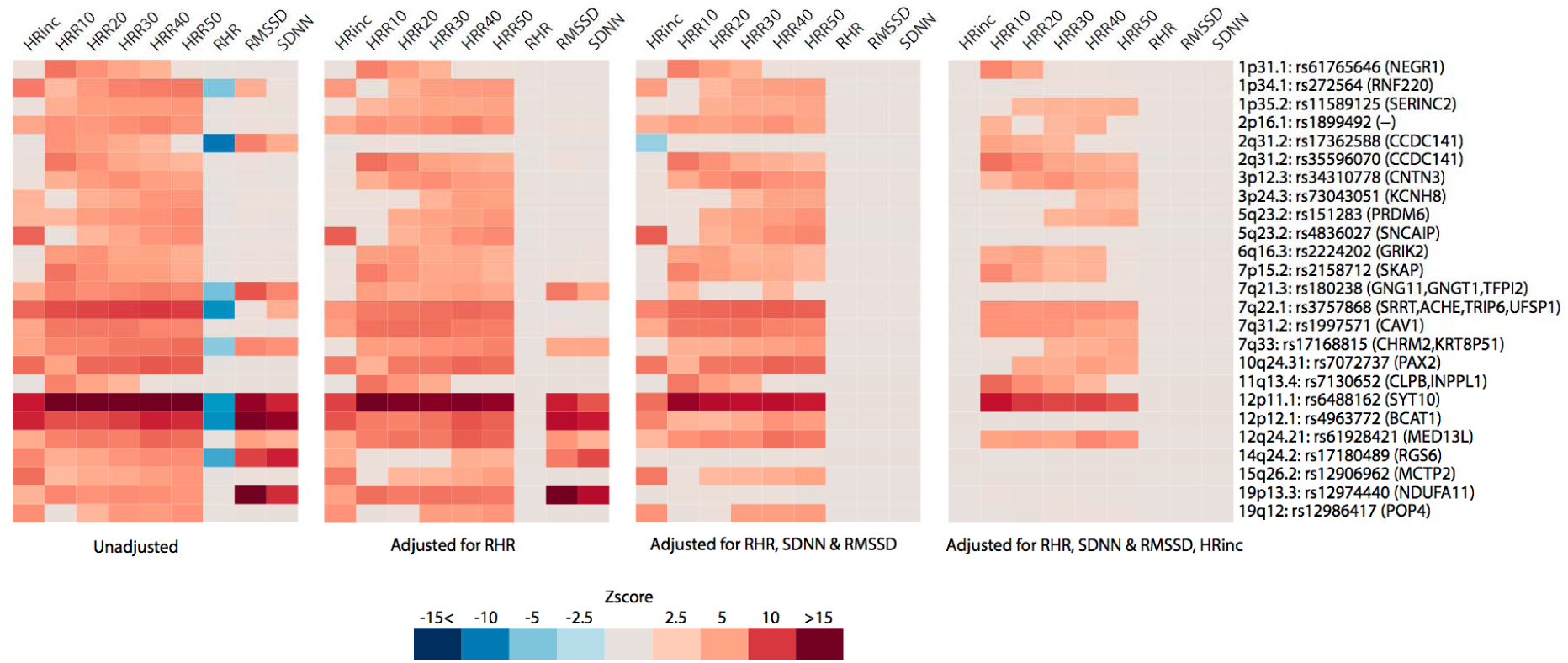
Pleiotropic effects of the 25 independent genetic signals on heart rate (HR) phenotypes. Four heat plots depict Z-scores of each SNP association with resting-HR (RHR), HR-variability (RMSSD and SDNN), HR-increase (HRinc) or HR-recovery (HRR10-50) in 1 univariate and 3 multivariable models (as described below each heat plot). Only Bonferonni P<0.05 significant associations are shown, Z-scores were aligned to the allele that increases HR-recovery. Nearby genes are shown between brackets.

To explore potential clinical relevancy, polygenic scores were constructed based on the genome wide significant SNPs. The primary outcome variable was parental age as proxy for cardiovascular- and all-cause mortality^14,46^. The choice of disease outcomes and phenotypes was based on the previous studies of HR-response to exercise in relation to ventricular arrhythmia (sudden death^4^), atrial fibrillation^47^, diabetes^48^, cancer^49^, or the importance of autonomic (dys)function in blood pressure^14^, reaction time, fluid intelligence^50^ and depression^51^. A higher polygenic score was consistently associated with an increased parental age of death (P=5.5×10^−4^). Upon further inspection, only a significant association was found with the father’s age of death (P=5.5×10^−4^ N=217,722), but not with the mother’s age at death (P=0.202, N=179,281). The association with increased parental lifespan may hint towards a potential association with all-cause mortality, which was not significant in UK Biobank (HR=0.924(0.055), P=0.186, N_cases_=10,717 (3.0%); cox survival model) but power is limited compared to parental age of death.

The polygenic score also strongly associated with lower diastolic blood pressure (P=2.0×10^−25^) and with lower odds of hypertension (P=2.3×10^−4^). The association with hypertension was dependent of diastolic blood pressure, as the association was abolished after introducing diastolic blood pressure into the model. We hypothesized that the strong association of the polygenic score with diastolic blood pressure may in fact be due to resting-HR because of its direct influence on diastolic blood pressure via peripheral resistance, considering the high genetic correlation of resting-HR with HR-increase and HR-recovery. After adjusting for resting-HR, the association with diastolic blood pressure was also abolished (P=0.126). No convincing associations were found between the polygenic score and atrial fibrillation, coronary artery disease, ventricular arrhythmia, diabetes or cancer. The results are shown in Table 2, **Supplementary Table 12** describes trait specific effects and **Supplementary Fig. 5** statistical power.

**Table 2.** The effect of the polygenic score of heart rate (HR) response to exercise on cardiovascular and non-cardiovascular phenotypes in the UK Biobank cohort, performed in participants that were not part of the discovery GWAS. Effect sizes are shown as the incremental change in phenotype for continuous phenotypes, or as odds ratio for binary traits, for one unit change in polygenic score. Every unit change in polygenic risk corresponds to one standard deviation change in HR-response to exercise. **Supplementary Table 12** shows the effect estimates per phenotype of HR-response.

| Trait or disease | Sample size<br>(% cases) | Effect size |  |  |
| --- | --- | --- | --- | --- |
|  |  | or odds<br>ratio | se / 95%CI | P |
| *Anthropometric* |  |  |  |  |
| Height (cm) | 420,910 | -0.1680 | 0.0612 | 0.006 |
| Weight (kg) | 420,697 | -0.0644 | 0.1361 | 0.636 |
| BMI (kg/m2) | 420,623 | 0.0326 | 0.0459 | 0.477 |
| *Cardiovascular risk factors* |  |  |  |  |
| DBP (mmHg) | 421,799 | -0.8240 | 0.0791 | 2.0×10<sup>-25</sup> |
| SBP (mmHg) | 421,797 | 0.0760 | 0.1560 | 0.626 |
| Pulse pressure | 421,797 | 0.9000 | 0.1140 | 3.0×10<sup>-15</sup> |
| Mean arterial pressure | 421,797 | -0.5240 | 0.0969 | 6.4×10<sup>-8</sup> |
| Hypertension | 422,334(33.85%) | 0.925 | 0.888-0.964 | 2.3×10<sup>-4</sup> |
| Coronary artery disease | 422,334(7.48%) | 1.022 | 0.950-1.100 | 0.554 |
| Atrial fibrillation | 422,334(3.71%) | 1.071 | 0.969-1.184 | 0.178 |
| Ventricular arrhythmia | 422,334(0.56%) | 0.868 | 0.674-1.117 | 0.271 |
| Diabetes Mellitus | 422,334(7.04%) | 1.072 | 0.996-1.155 | 0.064 |
| *Other* |  |  |  |  |
| Cancer (malignant) | 422,334(15.35%) | 0.983 | 0.932-1.036 | 0.512 |
| Depression | 422,334(14.35%) | 1.041 | 0.986-1.098 | 0.144 |
| Reaction time (ms) | 417,771 | -0.7016 | 1.0544 | 0.506 |
| Fluid intelligence score | 105,348 | -0.0645 | 0.0398 | 0.106 |
| Parental lifespan | 158,649 | 0.0792 | 0.0229 | 5.5×10<sup>-4</sup> |

## DISCUSSION

In this large-scale genetic study of HR-increase and HR-recovery in 58,818 participants, we identified 25 independent genome wide significant signals in 23 genetic loci. HR-increase and HR-recovery were found to be highly heritable and the majority of the loci were independently associated with HR-recovery. The polygenic score was not convincingly associated with mortality or disease.

The major finding was that a large proportion of candidate genes are involved in neuron biology, particularly at loci that are specific for HR-recovery. This, together with our pathway analyses, provides a new line of evidence that the autonomic nervous system is a major player in the regulation of HR recovery. Heart rate response to exercise, in particular HR-recovery, is largely dependent on parasympathetic reactivation and decrease of sympathetic activity in a gradual manner. These processes are orchestrated by neuronal signal transduction involving the brain (central command), periphery (chemoreflex, baroreflex, exercise pressor reflex), adrenal medulla and the actual nerves connecting them^15^.

The highest associated variant, rs6488162 in *SYT10,* encodes a Ca^2+^ sensor Synaptotagmin 10 that triggers IGF-1 exocytosis, protecting neurons from degeneration^52^. Other loci include the *ACHE* gene, the function of which can be strongly linked to neuronal function as it encodes the enzyme that catalyzes the breakdown of acetylcholine. *NEGR1*, Neuronal Growth Regulator 1, is essential for neuronal morphology. It has been shown by *in-vitro* and *in-vivo* experiments that *NEGR1* over- and under-expression is tightly associated with the number of synapses and proper development of neurite arborization and dendritic spines^53^. *GRIK2* (also named GluR6) encodes a subunit of a kainite glutamate receptor that is broadly expressed in the central nervous system where it plays a major role in nerve excitation^54^. *CHRM2* encodes M2 mAChR, which is the predominant form of muscarine cholinergic receptors in the heart. This gene fits very well with the specific association of rs17168815 (near CHRM2) with HR-recovery since this receptor specifically initiates negative chronotropic and inotropic effects upon binding with acetylcholine released by the postganglionic parasympathetic nerves, which are slowing down heart rate^55^. The gene *C19orf12* has an unknown function and is thought to encode a mitochondrial protein, there are several reports on mutations of *C19orf12* causing neurodegeneration^56^. The function of *MED13L* is also unclear, but is believed to encode a subunit that functions as a transcriptional coactivator for most RNA polymerase II-transcribed genes. *MED13L* knockdown in zebrafish causes abnormal effects on early migration of neural crest cells resulting in improper development of branchyal and pharyngeal arche, resembling key characteristics of *MED13L* mutations in humans^57^. *MED13L* mutations in humans are associated with intellectual disability, developmental delay and craniofacial anomalies, and resemble other common neurodevelopmental disorders^58^. *KCNH8* encodes a voltage gated potassium channel that is primarily expressed in components of the human central nervous system^59^ and is part of the Elk (ether-à-g-o go-like k) family of potassium channels that regulate neuronal excitation^59–61^. *CNTN3* (contactin-3) is a gene belonging to a group of glycosylphosphatidyl-anchored cell adhesion molecules that is thought to be closely involved in wiring of the nervous system and found predominantly in neurons^62,63^. In light of these findings, even *CCDC141* and not *TTN* (the main component of cardiac muscle), may be a plausible candidate gene, as it plays a crucial role during neuronal development^64^.

We observed that resting-HR, HR-variability, HR-recovery and HR-increase are highly correlated with each other on the genetic and phenotypic level. By jointly analyzing different HR-traits, instead of treating them as separate entities as has been done traditionally, it is possible to obtain additional insights into the mechanistic basis of HR phenotypes. On the phenotypic level, this helped us explain the strong association that was observed between the polygenic score and diastolic blood pressure; it was originating from resting-HR, which is more plausible since resting-HR is directly related to peripheral-resistance. On the genetic variant level, we observed a large proportion of genetic variants that are specifically associated with HR-recovery to contain neuronal genes as candidate genes. Previous GWAS studies of resting-HR have found genes predominantly enriched for terms related to cardiac structure^24^ and GWAS of HR-variability found genes involved in the sinoatrial node to be enriched^23^. In our analysis, the sinoatrial node genes *GNG11* and *RGS6* that were both previously associated with HR-variability^23^, were chiefly associated with HR-variability in this study as well. This emphasizes the importance for future follow-up studies to focus on extracting even more different HR phenotypes before, during and after exercise; so that they can be jointly analyzed to increase the resolution of SNP-phenotype of HR specific associations even further.

Observational studies have shown strong associations of HR-recovery and HR-increase with sudden cardiac death, all-cause death, cardiovascular death^4,5,46^ and even cancer^49^. These studies all suggested that autonomic impairment, the imbalance of vagal and adrenergic tone, increases the susceptibility to disease and mortality, and (although never shown) life-threatening arrhythmias. In this study, we observed that a genetically increased HR-recovery and HR-increase was significantly associated with higher parental age, but not with ventricular arrhythmia, atrial fibrillation or other diseases and phenotypes. Since the polygenic risk score was not significantly associated with the mother’s age of death, we could not reliably establish a true positive association with parental age. Although, the notion that life-threatening arrhythmia’s occur more often in men than in women could explain this discrepancy^65^. Regardless whether the association is a true-positive one, it is possible to conclude from our results that HR-response to exercise may not be as important for human life span compared to other more established risk factors like blood pressure, lipids, BMI or educational attainment^26^. The association with parental age should be followed-up in independent cohorts, but statistical power may be difficult realize given the exceptionally large sample size of this study. Future Mendelian randomization studies should be conducted in even larger cohorts and with other disease outcomes such as fatal arrhythmias, to provide a better understanding of the clinical consequences.

In conclusion, this is the first well-powered genetic study of HR-recovery and HR-increase, identifying 25 genetic signals in 23 loci to be genome wide associated. This study adds a new line of evidence for the fact that components of the autonomic nervous system are underlying inter-individual differences in HR-recovery.

## DISCLOSURES

The authors have declared that no conflict of interest exists.

### ACKNOWLEDGEMENTS

We would like to thank the Center for Information Technology of the University of Groningen for their support and for providing access to the Peregrine high performance computing cluster. We also thank dr. Thomas Teijeiro for his assistance with the Construe algorithm.

